# Comparison of morphological and physiological response to drought stress among temperate forest understory forbs and graminoids

**DOI:** 10.1101/2024.10.01.615773

**Authors:** Anja Petek-Petrik, Peter Petrík, Marika Halmová, Roman Plichta, Marie Matoušková, Kateřina Houšková, Markéta Chudomelová, Josef Urban, Radim Hedl

## Abstract

- Drought stress can profoundly affect plant growth and physiological vitality, yet there is a notable scarcity of controlled drought experiments focused on herbaceous species of the forest understory.
- In this study, we collected seeds from five forb and four graminoid species growing in the temperate forest understory of the Czech Republic. These seeds were germinated under controlled glasshouse conditions and subjected to moderate drought stress for five weeks. We assessed biomass partitioning, stomatal and leaf morphology, leaf gas exchange, minimum leaf conductance (*g*_min_), and chlorophyll fluorescence parameters.
- The comparison of two ecological guilds revealed that graminoids exhibited a higher root-to-shoot ratio, improved water-use efficiency, greater carboxylation efficiency, and enhanced non-photochemical quenching under drought conditions compared to forbs. In contrast, forbs had significantly lower *g*_min_, along with higher total biomass and total leaf area. Despite these differences in morpho-physiological functional traits, both groups experienced a similar relative reduction in biomass during drought stress. Key predictors of biomass accumulation under drought included photochemical quenching, stomatal traits, total leaf area and *g*_min_. A negative correlation between biomass and *g*_min_ suggests that plants with lower residual water losses after stomatal closure can accumulate more biomass under drought stress. Additionally, *g*_min_ was positively correlated with guard cell length, suggesting that larger stomata contribute to higher residual water loss.
- Graminoids exhibited morpho-physiological modifications that enhanced drought resistance, indicating a greater emphasis on stress tolerance as a survival strategy. In contrast, forbs maintained higher biomass and total leaf area, reflecting a competitive strategy for maximizing resource acquisition.

## 1. Introduction

Climate change is expected to exacerbate the impact of drought stress on forest understories, profoundly altering their structure and function (Koelemijer et al. 2022, Pozner et al. 2022, Tng et al. 2022, Deng et al. 2023). As global temperatures rise and precipitation patterns become more erratic, the frequency and severity of droughts are anticipated to increase (Cook et al. 2018, Schuldt et al. 2020, Pokhrel et al. 2021). The intensified drought stress can significantly reduce soil moisture levels, limiting water availability for understory vegetation, which typically comprises shrubs, young trees, and herbaceous plants. These understory species are crucial for biodiversity, providing habitat and food sources for a variety of wildlife (Cavard et al. 2011, Simonetti et al. 2013, Botequim et al. 2021). Herbaceous understory plants are also significant contributors to ecosystem carbon assimilation budget in natural forests (Petrík et al. 2024a). However, their mostly shallower root systems could make them particularly vulnerable to water scarcity (Canadell et al. 1996, McLachlan and Bazely 2001). Prolonged drought conditions can impair photosynthesis, decrease growth, lower reproduction ability and increase mortality rates among understory plants (Tng et al. 2022). Consequently, the resilience of forests understory to environmental stressors is compromised, potentially leading to long-term ecological imbalances and decreased forest health and productivity.

Plants exposed to drought stress close their stomata to prevent excessive water losses (Martin-StPaul et al. 2017, Petek-Petrik et al. 2023; Alongi et al. 2024). This also prevents CO_2_ intake which leads to a reduction of carbon assimilation via photosynthesis (Wilson et al. 2000). Lower assimilation negatively affects their carbon balance and less carbohydrates are available for growth (Attia et al. 2015). Drought stress also exuberates reactive oxygen species generation which can damage photosynthetic apparatus, further reducing the carbon assimilation potential of plants (Mukarram et al. 2021; Torun et al. 2024). Plants can partially mitigate these negative effects via changes in their morphological and physiological traits. Adjustment of morphological or physiological traits that has a positive impact on stress resistance is then called acclimation. For example, plants can change their carbon allocation patterns and invest more carbon into their root system rather than their aboveground foliage (Ye et al. 2021, Jiang et al. 2023, Reinelt et al. 2023). This increases their root-to-shoot ratio which enables them to access more water in the soil and reduce their total transpiration (Bacher et al. 2022). Moreover, the development of thicker and smaller leaves with lower specific leaf area (*SLA*) and higher leaf dry mass content (*LDMC*) correlates with higher osmotic potential of the leaves (Chelli et al. 2019, Blumenthal 2020, Bhusal et al. 2021) and higher water-use efficiency (*WUE*) (Horike et al. 2021, Zhong et al. 2022). Another potential acclimation response to drought stress can be the development of leaves with smaller stomata (Petrík et al. 2020, Petrík et al. 2022). Smaller stomata are faster in response to changing environmental conditions (Kardiman and Raebild 2018) and guard cell length (*GCL*) is negatively correlated with *WUE* (Petrík et al. 2023, Petrík et al. 2024b). The stomata size could potentially affect minimum leaf conductance (*g*_min_), which is residual water loss after stomatal closure through cuticle and incompletely closed stomata called ‘leaky stomata’ (Duursma et al. 2019). Recent studies have identified *g*_min_ as a critical trait for drought survival. Species exhibiting lower *g*_min_ are better able to maintain water status, thereby delaying hydraulic failure and reducing drought-induced mortality compared to species with higher *g*_min_ rates (Gleason et al. 2014, Duursma et al. 2019, Pereira et al. 2024). Several studies suggest that *g*_min_ may be plastic, with plants potentially acclimating to drought stress by reducing *g*_min_ (Le Provost 2013, Chen et al. 2020). In contrast, other research indicates that *g*_min_ is a conservative trait with low phenotypic plasticity and minimal acclimation potential (Schuster et 2017, Slot et al. 2021, Wang et al. 2024). Consequently, it remains an open question whether plants can modify *g*_min_ in response to drought stress. Furthermore, an increase in non-photochemical quenching as a relief pathway for excessive energy in chloroplasts can be seen as stress acclimation to protect photosynthetic apparatus from oxidative stress (Vastag et al. 2020, Turc et al. 2024). These acclimation strategies are crucial for maintaining the survival and ecological functions of understory herbs. However, the effectiveness of these responses might vary among ecological guilds and will be a determining factor in the resilience of forest ecosystems to the escalating impacts of climate change.

Forest understory graminoids and forbs exhibit distinct drought responses due to their differing morphological and physiological traits (Johnson et al. 2011, van Sundert et al. 2021). Graminoids, which include grasses and sedges, generally have narrow, linear leaves and high *WUE*, as their stomata tend to be smaller and more densely packed, reducing water loss through transpiration (Tobin et al. 1999, Hommel et al. 2014, Felsmann et al. 2018). They often have extensive fibrous root systems that allow for efficient water absorption from shallow soil layers (Šmilauerová and Šmilauer 2006). In contrast, forbs typically have larger leaf surfaces that can lead to higher water loss but also enable greater photosynthetic capacity under favourable conditions (Gobin et al. 2015, Felsmann et al. 2018). Forbs may develop deeper taproots, allowing access to deeper soil moisture, and can exhibit greater plasticity in their growth and phenology, adjusting their life cycles to periods of water availability (Zhang and Sun 2020). These differences result in graminoids often being more drought-resistant due to their conservative water use strategies, while forbs can be more drought-resilient, capable of rapid recovery following drought periods (Khan et al. 2022). This divergence in drought response strategies underscores the complex dynamics of forest understory ecosystems in the face of climate change.

Despite the growing recognition of climate change’s impact on forest ecosystems, there remains a significant lack of knowledge on the drought response of forest understory herbs. While substantial research has been conducted on the effects of drought on tree species and overall forest health, the responses of understory herbs—a critical component of forest biodiversity and function— are not well understood. This study seeks to compare the drought responses of native forbs and graminoids from temperate forest understories and to evaluate their acclimation potential. We anticipate that graminoids will exhibit greater resistance to drought stress compared to forbs, as evidenced by: (i) a lower relative reduction in biomass, (ii) smaller stomata, (iii) lower *SLA,* (iv) higher *WUE*, (v) higher non-photochemical quenching when exposed to drought stress. We also hypothesize (vi) that drought stress will induce reduction of *g*_min_ within forbs and graminoids groups.

## 2. Materials and methods

### 2.1 Plant material and seeds sampling site

The subjects of our study included nine herbaceous plant species: *Bromus benekenii* (Lange) Trimen, *Carex digitata* L., *Geum urbanum* L., *Impatiens parviflora* DC., *Lamium maculatum* (L.) L., *Melica nutans* L., *Poa nemoralis* L., *Veronica sublobata* M. A. Fisch., and *Viola mirabilis* L. These species are commonly found in temperate woodland understories of Central Europe and, except for the introduced *Impatiens parviflora*, form native plant communities in the region. Seeds were collected during the 2021 growing season in an oak-hornbeam forest located on the northeastern slopes of Děvín Hill (550 m a.s.l.), southeastern Czech Republic (48.873°N, 16.652°E). The site is prone to summer drought, with an average annual temperature of 9.6 °C and average annual precipitation of 524 mm.

### 2.2 Seedlings germination, experimental design and environment

The seeds collected in the field were initially subjected to a period of cold stratification at 1-3 °C for five weeks. Following cold stratification, seeds from the nine studied species were transferred to Petri dishes containing filter paper and 8 ml of distilled water. The Petri dishes were then placed in climate chambers (MLR-352H-PE) set at 70% relative humidity, with a photoperiod of 8h light at 30 °C, and 16h darkness at 20 °C.

Once germination was achieved, the seedlings were placed in a greenhouse (Mendel University, Brno, Czech Republic), from December 2021 to April 2022. The seedlings were potted into 36 plastic trays (40 × 30 × 7 cm) filled with 5 liters of a soil mixture (1:1 peat soil and sand) and set in a greenhouse maintained at 20 °C during the day (13h) and 12 °C at night (11h). The trays were organized into two experimental groups per species, each represented by four trays—two for control (well-watered) and two for drought treatment (water-deficit). Each tray contained 12 seedlings of one species, in total 48 individuals per species and 24 individuals per treatment.

Initially, all trays were irrigated with 400 ml of tap water every other day to maintain soil moisture conditions comparable to those in the Děvín forest. Following plant establishment, control treatment continued to receive 400 ml of tap water every other day for five weeks. In contrast, the drought treatment received approximately 150 ml of tap water every other day to induce water deficit conditions. The irrigation quantity was adjusted based on continuously measured soil moisture using the TMS-4 probes (TOMST, Prague, Czech Republic). The desired level of volumetric soil moisture was maintained at approximately 20% for control and 10% for drought treatments. This corresponded to –0.3 and –3 MPa of matric soil water potential, respectively. The volumetric soil moisture was recalculated into the soil metric water potential using the same substrate, 4 TMS probes and a WP4C Soil Water Potential Meter (Meter Group, Pullman, WA, USA). The temporal dynamics of soil water potential differed among control and drought and had similar dynamics among forbs and graminoids (Figure 1).

**Figure 1.**
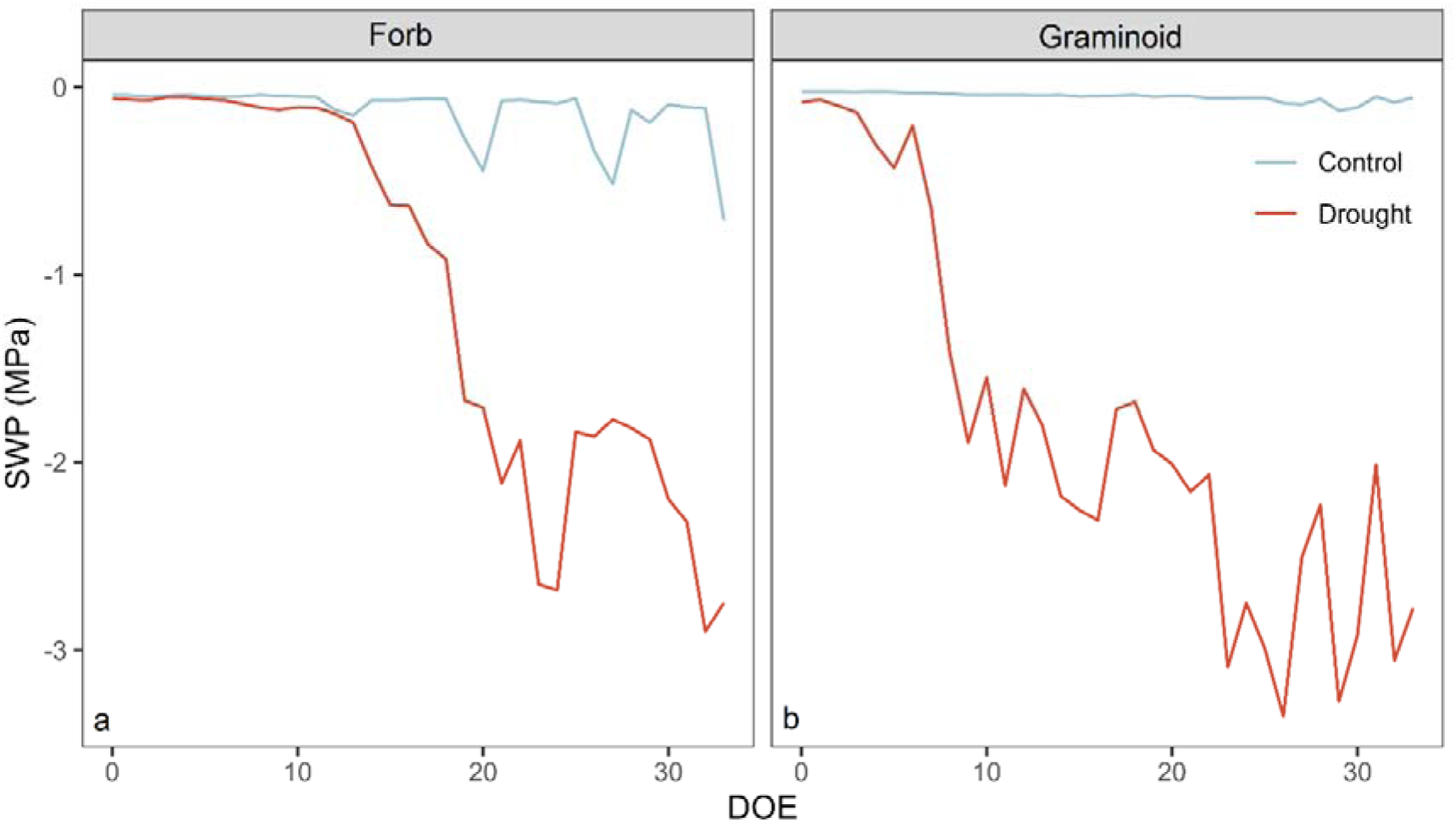
Temporal dynamics of soil water potential (SWP) plotted against the day of experiment (DOE) for control (blue) and drought (red) treatment. Daily averages are presented for forbs (a) and graminoids (b).

### 2.3 Leaf morphology and biomass

After the five-week experimental period, individuals from the control and drought treatment groups were harvested for biomass and leaf morphological analyses. Individuals from each species and treatment group were carefully excavated from the soil and subsequently dissected into leaf, shoot, and root components. Before further analysis, soil was removed from leaves, shoots, and roots using a fine brush. The dissected parts of each individual were weighed with an analytical balance (KERN EWB 620-2M) to determine their biomass. All leaves from each individual were scanned using a standard flatbed scanner (Epson DS-1630, 300 dpi), and the total leaf area (*TLA*) was determined using ImageJ software (National Institute of Health, USA). Previously scanned leaves were oven-dried at 60 °C for 48h and weighed using an analytical balance (KERN EWB 620-2M) to obtain dry biomass. These measurements were used to determine the specific leaf area (*SLA*), calculated as the ratio of *TLA* to leaf dry weight. The shoot and leaf biomass of each individual were summed to obtain the aboveground biomass (*AGB*). The root-to-shoot ratio (*R:S*) was subsequently calculated as the ratio of belowground biomass (*BGB*) to *AGB*. A total biomass (*B*) was calculated as the combined sum of *AGB* and *BGB*. For *SLA* and *LDMC*, the sample size ranged from 12 to 24 individuals, while biomass metrics were assessed on 15 to 24 individuals per species and treatment.

### 2.4 Stomatal morphology

The stomatal imprints were collected at the end of the five-week drought stress period. The stomatal imprints were taken from fully developed apical leaves using the ‘kollodium method’ (Petrík et al. 2020), in which transparent nail polish was applied to the abaxial and adaxial sides of the leaves and transferred to a microscope slide using transparent tape. The imprints were used for capturing of digital photographs with the use of a Levenhuk MED 30T (Levenhuk, Tampa, USA) microscope equipped with Delta Optical DLT-Cam Pro 12MPx (Delta Optical, Poland). One photograph of imprint from each leaf and side was taken at 20 x 10 magnification. The stomatal morphological traits were further analysed with ImageJ software. The length of guard cell (*GCL*) was measured on three randomly selected stomatal cells per photograph and then averaged per individual. The stomatal density (*SD*) was assessed as the total number of stomata per photograph (0.0415 mm^2^), recalculated to *SD* per 1 mm^2^. The sample size for both *GCL* and *SD* ranged between 10 to 24 individuals per species and treatment. Collecting stomatal imprints was not feasible for some drought-stressed individuals due to their leaves being too small and wilted.

### 2.5 Minimum leaf conductance

The minimum leaf conductance (*g*_min_, mmol m^-2^ s^-1^) was determined on five to six detached leaves for each species and treatment, by the “mass loss of detached leaves’’ method (Blackman et al. 2019, Duursma et al. 2019, Petek-Petrik et al. 2023). This method involves monitoring branch weight over time under stable atmospheric conditions as the leaves desiccate. Leaves from the plants were excised and left to saturate in water overnight. The cut was isolated with the tape, leaving the leaf transpiring from both sides, while attached within a controlled-climate chamber (Fytoscope FS 130, Photon Systems Instruments, Drásov, Czech Republic) to dry out. Within the chamber, relative humidity was maintained at 60% and temperature was set at 20 °C. The climate chamber was placed in an air-conditioned room with a temperature set at 20 °C to avoid rapid changes in the conditions when opening the chamber to conduct the measurements. The loss of leaf mass was measured with an analytical balance with a scale resolution of 0.1 mg (VWR TA314i, Leuven, Belgium) every 5-10 minutes for the first hour, and then every 30 minutes thereafter, until 20 data points were obtained. The *g*_min_ was then calculated from the slope of the linear part of the leaf mass loss (mmol) over time (seconds), as follows: *g*_min_ = (slope * (atmospheric pressure / vapour pressure deficit)) / leaf area. Following the measurement of *g*_min_, the projected leaf area was determined using scans (Epson, DS-1630) and analysed with Image J software to recalculate the *g*_min_ values per square meter.

### 2.6 Gas-exchange measurements and fluorescence measurements

The light-saturated net assimilation rate (*A*), stomatal conductance (*g*_s_), transpiration (*E*) and the intercellular concentration of CO_2_ in the leaves (*C*_i_) were measured by the infrared gas analyser (LI-6800, LI-COR Inc., Lincoln, Nebraska, USA). Intrinsic water use efficiency (*WUE*i) and the instantaneous carboxylation efficiency (*A/C*_i_) were calculated as the ratio of *A* and *g*_s_ and *A* and *C*_i_, respectively. All measured leaves were enclosed into the chamber and exposed to the following conditions: the airflow rate of 600 μmol s^-1^, the CO_2_ concentration of 400 μmol CO_2_ mol^-1^, relative humidity of 50-60% and the saturated photosynthetic photon flux density (PPFD) of 1000 μmol photons m^-2^ s^-1^. The leaf area enclosed into the chamber was 2 cm^2^. Measurements were carried out randomly on 4 to 24 individuals per species and treatment the week preceding biomass sampling. Due to the small and fragile leaves, gas-exchange measurements could not be obtained for drought-stressed *Bromus benekenii* individuals.

The open state quantum efficiency of photosystem II (*Fv’/Fm’*), photochemical quenching (*qP*) and non-photochemical quenching (*qN*) were also measured by infrared gas analyser LI-6800 (LI-COR Inc., Lincoln, Nebraska, USA) using rectangular multiphase flash (MPF) during the same measurement days as gas-exchange measurements, on the same leaves. MPF was set to a margin of 5 points, an outrate of 100 data points per second, a maximal saturating pulse during the flash of 10 194 μmol photons m^-2^ s^-1^ and modulation rate of 250 000 Hz. The sample size of the fluorescence measurements corresponded to the gas-exchange measurements and ranged from 4 to 24 individuals per species and treatment.

### 2.7 Statistical analysis

All statistical analyses were conducted using the R 4.2.1 software (R Core Team, Vienna, Austria). The species-level averages of all measured morphological and physiological traits under control and drought treatments, along with 95% confidence intervals, are shown in the Appendix (Table A1). As the sample size and variance between the groups were not homogeneous, we applied the Kruskal-Wallis test and Dunn’s non-parametric post-hoc test to analyse the differences between the treatments and species (Appendix Table A2; Figure A1, A2). We conducted the individual-level Pearson correlation analysis between the traits separately for control and drought treatment (Appendix Figure A3, A4) by corrmorant package (Link 2020).

The species were further divided into two ecological guild groups for further analysis. The forbs group included *Geum urbanum*, *Impatiens parviflora*, *Lamium maculatum*, *Veronica sublobata* and *Viola mirabilis,* while the graminoid group comprised *Bromus benekenii*, *Carex digitata*, *Melica nutans* and *Poa nemoralis*. Differences between the ecological guild groups and treatments were analysed using the Kruskal-Wallis test (Appendix Table A2) and Dunn’s non-parametric post-hoc test (Figure 2, 3). Coordination between traits and the distribution of ecological guild groups and control-drought treatments was tested by principal component analysis (PCA) at the individual level (Figure 4). The representation of traits within the first three principal components of the PCA system was tested by cumulative Cos2 of the variables by factoextra package (Kassambara and Mundt 2020). The relationship between selected traits at the individual level was tested by linear and logarithmic regression separately for the control and drought treatment across all species (Figure 5).

**Figure 2.**
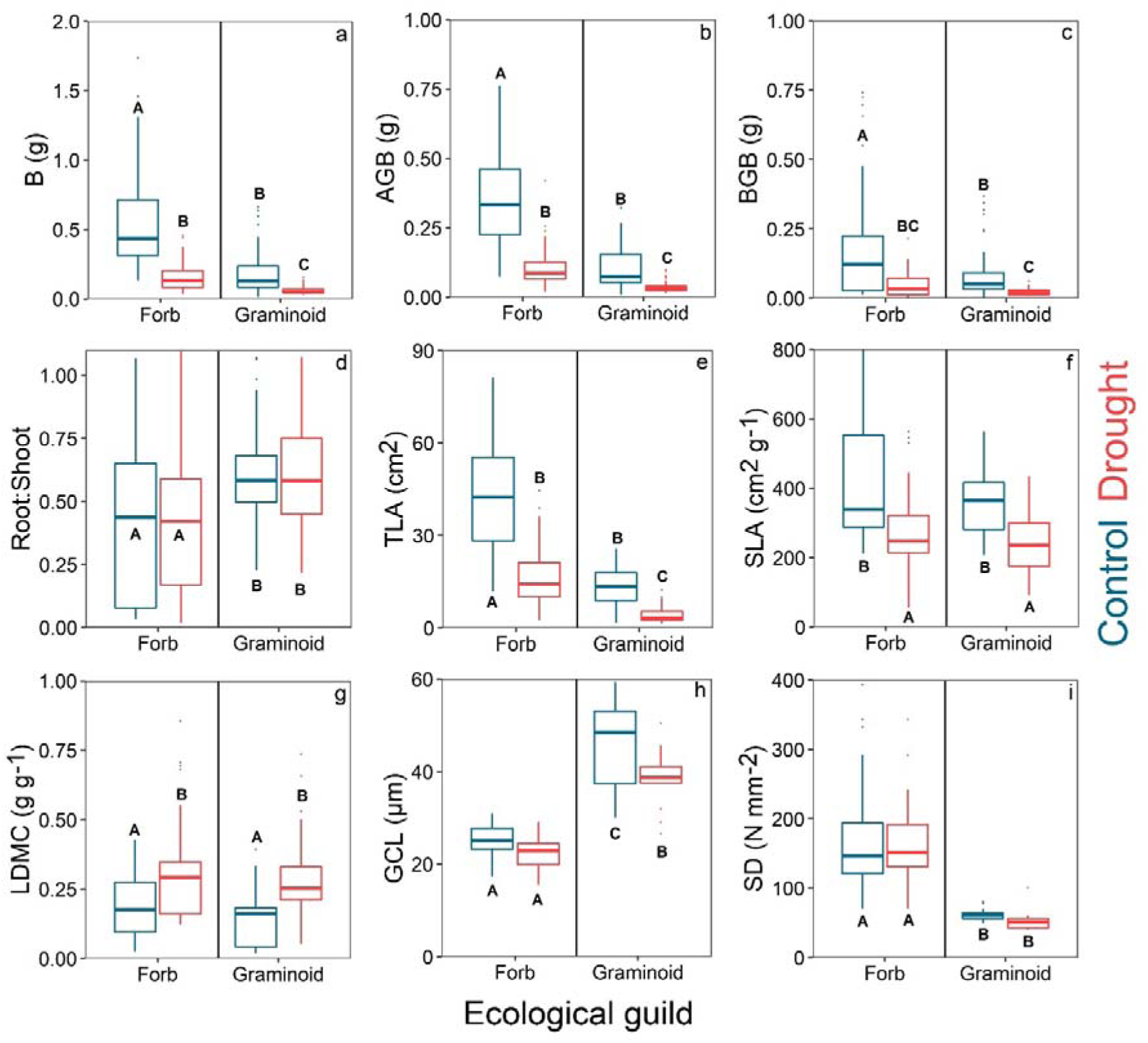
Ecological guild level boxplots of total biomass (a), aboveground biomass (b), belowground biomass (c), root:shoot ratio (d), total leaf area (e), specific leaf area (f), leaf dry matter content (g), stomatal guard cell length (h) and stomatal density (i).

## 3. Results

### 3.1 Morphological response to drought stress

The drought stress had a significant impact on biomass and leaf morphology, but a limited impact on stomatal morphology in both forbs and graminoids. The Kruskal-Wallis test revealed that ecological guild, drought treatment and their interaction had a significant effect on total biomass (*B*), aboveground biomass (*AGB*), belowground biomass (*BGB*) and total leaf area (*TLA*) of the plants (Appendix Table A2). Drought stress imposed a reduction in all three-biomass metrics and *TLA* for both ecological guilds (Figure 2a, b, c, e). The absolute reduction in these metrics was greater for forbs than graminoids; however, the relative reduction was similar, with *B* reduced by 28% for forbs and 31% for graminoids. The biomass was consistently reduced between above- and below-ground parts, resulting in the similar root-to-shoot ratio (*R:S*) of control and drought stress groups. Stomatal density (*SD*) was significantly higher for forbs, while *R:S* was significantly higher for graminoids. However, drought stress did not alter these traits (Figure 2d, i). Specific leaf area (*SLA*) and leaf dry matter content (*LDMC*) did not differ between the forbs and graminoids, and drought stress caused a similar reduction in *SLA* and increase in *LDMC* in both groups (Figure 2f, g). Graminoids exhibited significantly higher length of guard cell (*GCL*) compared to forbs. Drought stress significantly reduced *GCL* in graminoids but had no effect on forbs (Figure 2h). The species-specific averages and variability intervals, together with Dunn’s post-hoc tests, can be found in the Appendix (Table A1; Appendix Figure A1, A2).

### 3.2 Physiological response to water-deficit

Drought stress slightly decreased intrinsic water use efficiency (*WUEi*) in forbs and reduced assimilation, transpiration, and photosynthetic efficiency in both forbs and graminoids. Non-parametric tests revealed that drought treatment significantly affected light-saturated net assimilation rate (*A*), transpiration (*E*), and stomatal conductance (*g*_s_) in both plant groups (Figure 3a, b, c). Whole-plant transpiration (*E*TLA*) was 80.51 mmol s^-1^ (±21.34SE) for control forbs, 3.17 mmol s^-1^ (±0.73SE) for droughted forbs, 20.52 mmol s^-1^ (±8.33SE) for control graminoids and 0.77 mmol s^-1^ (±0.38SE) for droughted graminoids. The reduction in *A* was notably more pronounced in forbs compared to graminoids, resulting in a significant decrease in *WUEi* for forbs but not for graminoids (Figure 3d). Additionally, the instantaneous carboxylation efficiency (*A/Ci*) under drought condition was significantly higher for graminoids compared to forbs (Figure 3e). Although drought treatment did not significantly affect minimum leaf conductance (*g*_min_) in either group, graminoids exhibited higher *g*_min_ than forbs under drought conditions (Figure 3f). Our results therefore also support the findings by Schuster et al. 2017 that plants do not alter their *g*_min_ because of low phenotypic plasticity of cuticular water permeability. Drought treatment significantly reduced the open state quantum efficiency of photosystem II (*Fv’/Fm’*) in both groups (Figure 3g). Graminoids consistently showed lower *Fv’/Fm*’ values compared to forbs in both control and drought treatments. The photochemical quenching (*qP*) was reduced in forbs under stress, whereas no significant change was observed in graminoids (Figure 3h). Conversely, drought treatment had a positive effect on non-photochemical quenching (*qN*) in both forbs and graminoids (Figure 3i), with graminoids exhibiting higher *qN* values than forbs in both control and drought conditions.

**Figure 3.**
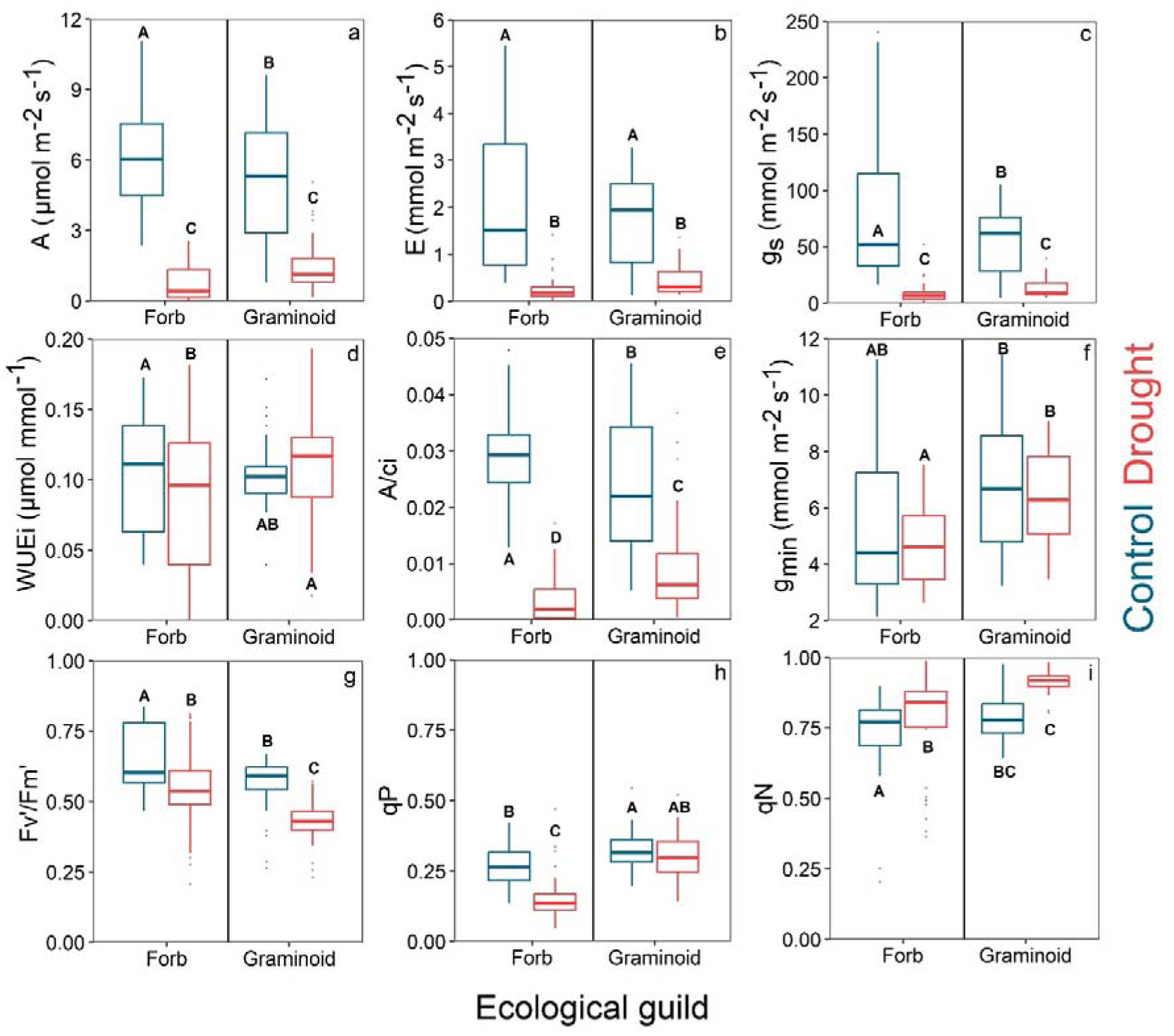
Ecological guild level boxplots of assimilation rate (a), transpiration rate (b), stomatal conductance (c), water use efficiency (d), carboxylation efficiency (e), minimal leaf conductance (f), open state quantum efficiency (g), photochemical quenching (h), non-photochemical quenching (i).

### 3.3 Coordination and trade-offs between traits under drought stress

Drought stress led to reductions in biomass, leaf morphological adjustment, and increased stomatal and non-stomatal limitation of assimilation in both forbs and graminoids (Figure 4a, c). The differences in stomatal morphological adjustment, *qP* and *qN* responses to drought stress among forbs and graminoids resulted in a clear separation between the groups in the PCA (Figure 4b, d). The first three principal components of the PCA explain all parameters well, except *R:S* and *WUEi* (Figure 4e). The low representation of *R:S* and *WUEi* (low cos2 values) is due to their high variability within groups and a weaker influence of drought stress (*R:S* showed no change and only forbs differed in *WUE*i). The PCA revealed coordination between plant morphological and physiological traits, with biomass traits (*B*, *AGB*, *BGB*) positively correlated with *TLA*, *SD*, gas-exchange parameters (*A, E, g*_s_, *A/Ci*) and *Fv’/Fm’*, while negatively correlated with *GCL*, *qN* and *LDMC* (Figure 4a, c).

**Figure 4.**
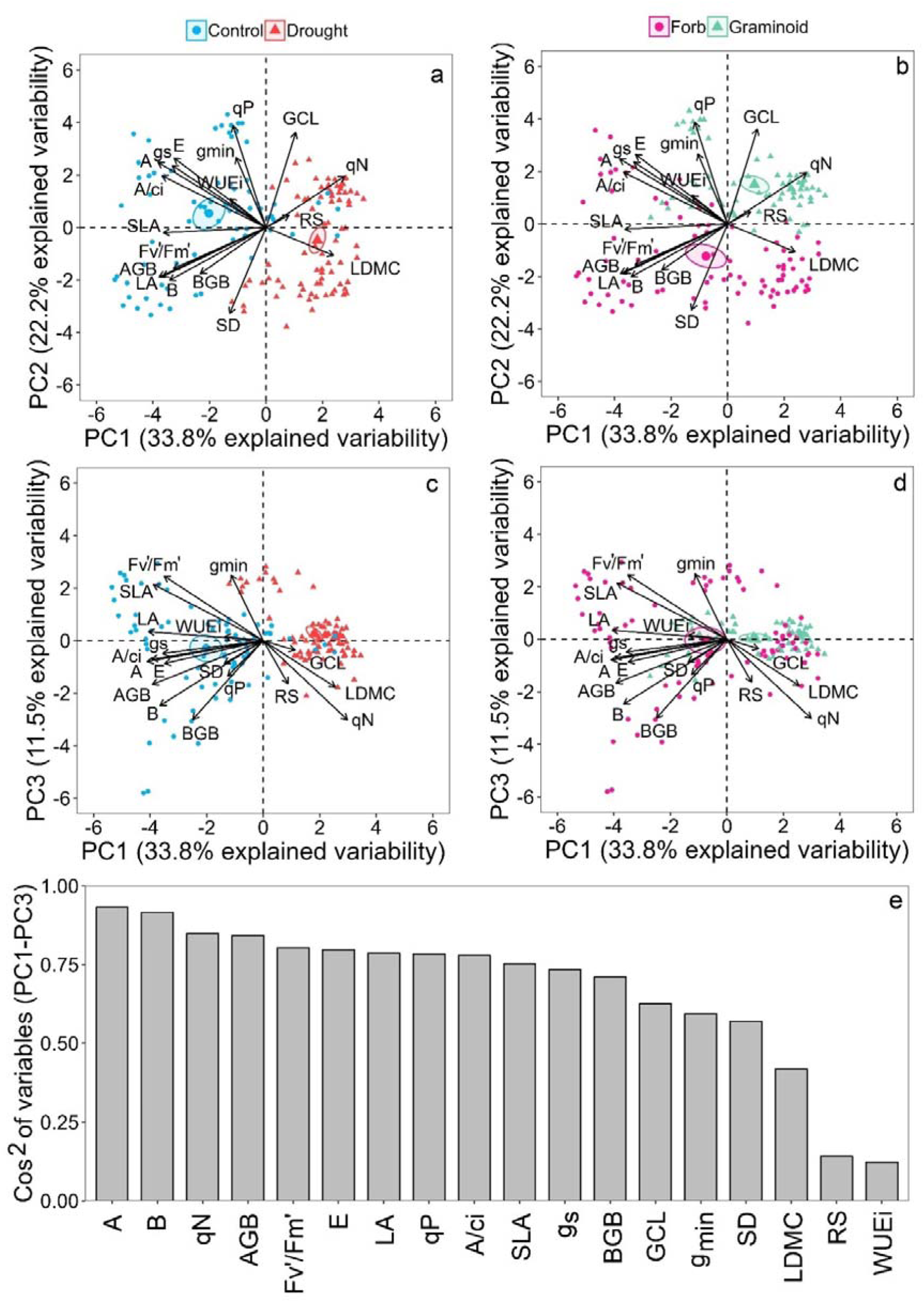
Principal component analysis biplots for the first three principal components illustrating differentiation between control and drought treatments (a,c) and between ecological guilds (b,d). The square cosines for each variable across the three principal components display the representation of the variables in the PCA (e). For the explanation of variables’ abbreviations see Figures 2 and 3.

To further analyse the data, we performed separate correlation analysis for the control and drought treatment including both ecological guilts (Appendix Figure A3, A4). The correlation matrix for the control group exhibited generally lower correlation strength and fewer significant relationships (24 at *p* < 0.05) between traits compared to the drought group (40 at *p* < 0.05). Specifically, *B* negatively correlated with *g*_min_ under drought conditions while not under control conditions (Figure 5a). Additionally, *g*_min_ showed a positive correlation with *GCL* and a negative correlation with *SD* under drought (Figure 5b, c). The negative relationship between *GCL* and *SD* represents a trade-off due to spatial constraints on the leaf (Figure 5d). Therefore, plants with lower *g*_min_, lower *GCL* and higher *SD* accumulated more *B* under drought stress (Figure 5, Appendix Figure A4).

**Figure 5.**
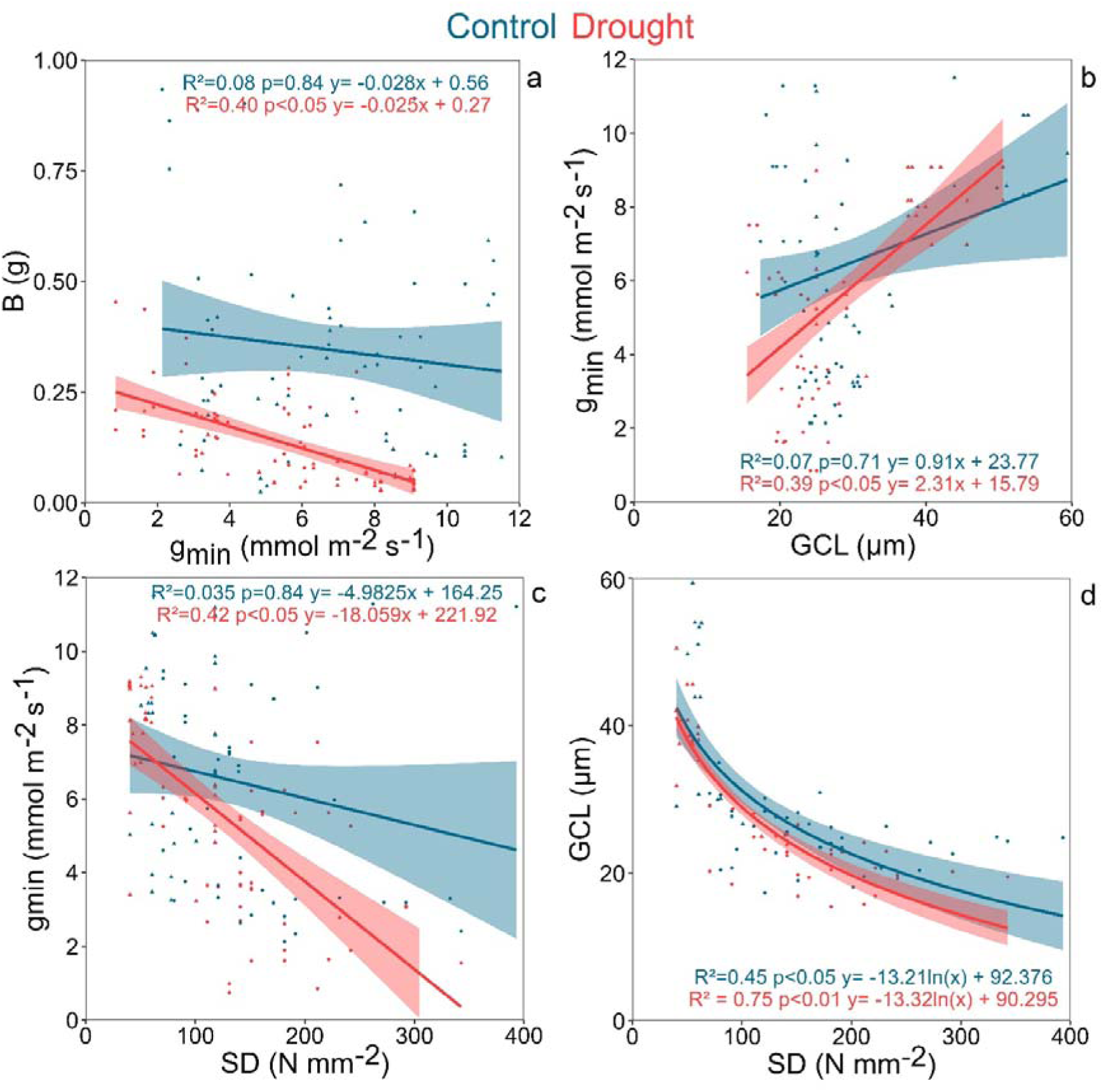
The linear relationships between biomass and minimal leaf conductance (a), guard cell length and minimal leaf conductance (b), stomatal density and minimal leaf conductance (c) and the logarithmic relationship between stomatal density and guard cell length (d), for both control (blue) and drought (red) treatments.

## 4. Discussion

Experimental studies on drought stress in native forest understory herbaceous species under controlled conditions are exceedingly scarce. Our research revealed significant differences in drought responses between forest understory forbs and graminoids, with graminoids exhibiting a greater capacity for acclimation to drought stress compared to forbs. Additionally, we elucidated the relationships between functional morphological and physiological traits, which are potentially critical for assessing drought resistance in forest understory herbaceous species.

### 4.1 Drought stress impact on morphological traits

Exposure to drought stress affected biomass accumulation, leaf and stomatal morphological traits, with the impact varying according to the ecological guild (Figure 2). Reduced water availability commonly leads to biomass reduction by limiting photosynthetic assimilation through stomatal closure, which in turn constrains growth (Flexas et al. 2008, Pirasteh-Anosheh et al. 2016). Under well-watered conditions, forbs generally exhibited higher aboveground and belowground biomass compared to graminoids (Grime and Hunt 1975). Graminoids typically possess extensive, shallow, yet widely spread fibrous root systems (Bardgett et al. 2014), whereas forbs often have deeper taproots or a combination of taproots and fibrous roots, allowing them to access moisture from deeper soil layers (Nippert and Knapp 2007). Despite this, no significant differences in belowground biomass were observed between forbs and graminoids under drought stress. On the other side, consistent with the findings of Zwicke et al. (2016), we observed that graminoids maintained a higher root-to-shoot ratio (*R:S*) under drought stress compared to forbs, though it is worth to note that drought stress had no effect on *R:S* either in forbs nor in graminoids. A higher *R:S* typically indicates that plants can access more water relative to their transpiring surface area (Bacher et al. 2022). Consequently, the elevated *R:S* observed in graminoids may enhance their drought resistance relative to forbs. In our experiment, the soil trays were shallow, which may have constrained the development of deeper rooting systems in the herbaceous plants. Therefore, our results likely reflect conditions similar to those encountered in shallow forest soils.

The relative reductions of the total leaf area (*TLA*), specific leaf area (*SLA*) and increase in leaf dry matter content (*LDMC*) are comparable between the forbs and graminoids. Although the reduction of *TLA* is typically associated with decreased assimilation and growth, it can also be interpreted as an acclimation response aimed at reducing overall transpiration and thereby slowing soil water depletion. *SLA* is a functional trait often used to characterise the resource acquisition strategy but also the potential stress tolerance of plants (Maire et al. 2015). Low *SLA* and high *LDMC* are generally considered as drought adaptation as they correlate with a higher osmotic potential of leaves (Blumenthal 2020, Bhusal et al. 2021). Graminoids have narrow leaves with a high surface-area-to-volume ratio, which helps reduce water loss through transpiration (Givnish 1978). This adaptation is particularly advantageous in dry or variable environments, allowing graminoids to conserve water and maintain hydration in conditions where it might be limited (Larcher 2003). In contrast, forbs have broader leaves that, while potentially increasing transpiration rates and water loss, also facilitate greater photosynthetic activity when water availability is available (Sage and Monson 1999). Consequently, the lower *TLA* observed in graminoids results in reduced whole-plant transpiration compared to forbs, leading to slower soil water depletion.

Forbs exhibited significantly shorter guard cell length (*GCL*), and higher stomatal density (*SD*) compared to graminoids. Additionally, drought treatment resulted in a reduction of *GCL* in graminoids, but not in forbs. This indicates that graminoids may possess greater phenotypic plasticity in their stomatal traits compared to forbs (Frank and Beerling 2009). Both *GCL* and *SD* can substantially influence stomatal conductance (*g*_s_) and the overall stomatal responsiveness of plants (Harrison et al. 2020, Murray et al. 2020, Gibbs et al. 2021). Furthermore, *GCL* has been shown to negatively correlate with water use efficiency (*WUE*), suggesting that plants with smaller stomata are better able to sustain carbon assimilation while conserving water (Petrík et al. 2023, Petrík et al. 2024). Stomatal morphology exhibits a degree of plasticity and can vary from year to year (Petrík et al. 2022). The observed reduction in *GCL* for graminoids in our study can be interpreted as a drought acclimation response (Liu et al. 2023). In contrast, the absence of *GCL* adjustment in forbs suggests a stronger genetic control of stomatal traits. These findings imply that graminoids exhibit greater resistance to drought stress in terms of growth and biomass allometry compared to forbs. Additionally, graminoids demonstrate higher plasticity in stomatal morphology, potentially enabling more effective acclimation to drought stress.

### 4.2 Physiological response to drought stress

Drought stress caused reduction of assimilation, transpiration and stomatal conductance in both forbs and graminoids (Figure 3). The relative reduction of these fluxes was greater for forbs, than for graminoids, parallel to differences in biomass accumulation. Forbs also experienced significant reduction of intrinsic WUE (*WUE*i) under drought treatment, compared to control. The *WUE*i of graminoids under drought treatment was higher than control, but not significantly. Conversely, the *WUE*i of drought stressed graminoids was significantly higher than *WUE*i of drought stressed forbs. Generally, high *WUE*i in grasses (Taylor et al. 2010) enables them to accumulate more carbon and also results in higher reproductive capacity compared to forbs under drought conditions. Graminoids exposed to drought stress also had significantly higher instanenous carboxylation efficiency (*A/Ci*) compared to drought stressed forbs. This means that graminoids were able to better mitigate non-stomatal limitations of photosynthesis like Rubisco efficiency or electron transport rate (Alongi et al. 2024). Overall, graminoids showed greater resistance (lower relative reduction) of their assimilation and stomatal conductance than forbs. Similarly, research by Griffin-Nolan et al. (2019) found that in drought-stressed environments, graminoids generally experience a smaller reduction in these physiological parameters than forbs, which contributes to their greater drought resistance. Moreover, Lawson and Vialet-Chabrand (2018) highlighted that graminoids are better adapted to fluctuating water availability, partly due to their ability to modulate stomatal conductance more effectively than forbs. Furthermore, significantly higher *WUE*i and *A/Ci* of graminoids under drought stress suggests they can utilize the water and carbon for assimilation more effectively than forbs. These findings underscore the functional differences between plant types and their strategies for coping with environmental stress.

The different strategy of forbs in coping with drought stress could also be seen in the residual water loss from leaves, characterised as the minimum leaf conductance (*g*_min_). This trait could play a decisive role in surviving the critical lack of water in soil (Duursma et al. 2019). On one side we observed no significant changes in *g*_min_ due to drought exposure for either forbs or graminoids. This supports the current understanding that *g*_min_ is a very conservative trait with limited phenotypic plasticity (Slot et al. 2021, Wang et al. 2024). Nevertheless, forbs showed significantly lower *g*_min_ under drought stress compared to graminoids. Specifically, forbs have 50% lower *g*_min_ than graminoids but possess 57% greater *TLA*. Consequently, the total residual water losses (*g*_min_ * *TLA*) are comparable between the two drought-stressed groups. Therefore, forbs and grasses employ different strategies to achieve similar total residual water losses at the whole plant level.

Both graminoids and forbs also experienced non-stomatal limitation of photosynthesis apparent from reduction of open state quantum efficiency (*Fv‘/Fm‘*) under drought stress. The reduction of *Fv‘/Fm‘* under drought stress is usually caused by oxidative damage of photosystem II (Colom and Vazzana 2003, Konôpková et al. 2018). Graminoids showed significantly lower *Fv‘/Fm‘* under drought stress compared to forbs. On the other hand, the non-photochemical quenching (*qN*) of graminoids was significantly higher than forbs under drought stress. Therefore, graminoids reduced their photochemical energy pathway in order to increase the *qN* and increase their photoprotection ability (Turc et al. 2024). There is also some evidence that increasing *qN* can lead to higher stomatal sensitivity and therefore faster stomatal closure, which can have also positive impact on water retention of the plants (Glowacka et al. 2018). The significantly higher *qN* in graminoids compared to forbs may enable them to dissipate excessive energy more efficiently and better protect their photosynthetic apparatus under drought stress.

### 4.3 Traits correlation

The PCA analysis provided a clear morpho-physiological division between the control vs. drought treatment and graminoids vs. forbs (Figure 4). The analysis also revealed both coordination and trade-offs between the traits. A more detailed look into the relationships between the traits was then achieved with separate correlation analyses for control and drought treatments (Appendix Figure A3, A4). The photosynthetic trait *Fv‘/Fm‘* showed the strongest correlation with the biomass traits *AGB*, *B* and *BGB*, compared to *A* or *A/Ci*. Moreover, *SLA* was also positively correlated with the *Fv‘/Fm‘* and biomass traits, confirming the general theorem of *SLA* being a functional trait reflecting the resource acquisition strategy of plants (Wilson et al. 1999, Maire et al. 2015). The strong correlation between *Fv‘/Fm‘*, biomass traits, and *SLA* suggests that the non-stomatal limitation explains better biomass accumulation differences under drought stress than stomatal limitation effects (*A*, *A/Ci*).

An additional important trait coordinated with biomass accumulation under drought was *g*_min_. We found a negative significant relationship between *g*_min_ and *AGB*, *BGB* and *B*, therefore individuals with lower *g*_min_ were able to accumulate more biomass under drought than individuals with higher *g*_min_ (Figure 5a). Moreover, we found that *GCL* has a positive and *SD* negative impact on *g*_min_ of the plants under drought stress (Figure 5b, c). This suggests that not only cuticle, but also stomata are significantly affecting the total *g*_min_ (Duursma et al. 2018), as it could be assumed that probability of not-completely closed stomata (so called ‘leaky stomata’) is increasing with the increasing *SD*. Leaky stomata accounted for 35% of *g*_min_ in *Hedera helix* (Šantrůček et al. 2004) and for 50-94% *g*_min_ in coniferous tree species (Brodribb et al. 2014). While we cannot precisely quantify how much of *g*_min_ occurs through stomata versus the cuticle, the explanatory power of 39% (*GCL*) and 42% (*SD*) in linear regression with *g*_min_ theoretically corresponds to the value found experimentally by Šantrůček et al. (2004). The *g*_min_ is a critical trait to predict the drought mortality timing in plants, as higher *g*_min_ means faster depletion of water reserves after stomatal closure (Petek-Petrik et al. 2023, Waite et al. 2024, Ziegler et al. 2024). Our study suggests that *g*_min_ is also an important trait for biomass accumulation under drought stress, with plants showing lower *g*_min_ likely accumulating higher biomass due to better water retention capacity.

## 5. Conclusion

Drought differentially influenced the morphological and physiological traits of temperate forest forbs and graminoids. Our study indicates that graminoid species demonstrate greater drought resistance than forb species. In contrast, forbs displayed a higher competitiveness, characterized by the ability to maintain higher total leaf area and biomass. Additionally, our findings reveal that minimum leaf conductance plays a significant role in biomass accumulation during drought stress. Specifically, individuals exhibiting lower minimum leaf conductance were able to accumulate greater aboveground, belowground, and total biomass compared to those with higher minimum leaf conductance. This relationship emphasizes the significance of water retention mechanisms in improving drought resilience. Overall, these findings highlight the distinct adaptive strategies of forbs and graminoids in response to drought.

## Acknowledgements

We thank Daniel Kurjak for providing the microscope used to capture photographs of the stomatal imprints and MENDELU for providing the greenhouse. This research was funded by project 21-11487S of the Czech Science Foundation. MH, RH and MC were supported by the long-term research development project RVO 67985939 to the Institute of Botany of the Czech Academy of Sciences. RP was supported by the project INTER-TRANSFER LTT20017 of the Ministry of Education, Youth and Sports of the Czech Republic. MH was supported by the internal project of Palacký University PrF-2024-001.

## Authors contributions

RH and JU conceived the research idea. RH, APP, RP, and JU designed the methodology. APP, PP, MK, JU, and MM carried out the measurements in the greenhouse and laboratory. MK and MC collected the seeds in the field. KH carried out the seedling germination. PP and RP analysed the data. APP and PP led the writing of the manuscript. All authors contributed critically to the drafts and gave final approval for publication.

## List with abbreviations

| TRAITS | ABBREVIATIONS | UNITS |
| --- | --- | --- |
| TOTAL BIOMASS | *B* | g |
| ABOVEGROUND BIOMASS | *AGB* | g |
| BELOWGROUND BIOMASS | *BGB* | g |
| TOTAL LEAF AREA | *TLA* | cm<sup>2</sup> |
| ROOT-TO-SHOOT RATIO | *R:S* | - |
| SPECIFIC LEAF AREA | *SLA* | cm<sup>2</sup> g<sup>-1</sup> |
| LEAF DRY MATTER CONTENT | *LDMC* | g g<sup>-1</sup> |
| STOMATAL DENSITY | *SD* | N mm<sup>-2</sup> |
| LENGTH OF GUARD CELL | *GCL* | μm |
| INTRINSIC WATER USE EFFICIENCY | *WUEi* | μmol mmol<sup>-1</sup> |
| ASSIMILATION RATE | *A* | μmol m<sup>-2</sup> s<sup>-1</sup> |
| TRANSPIRATION RATE | *E* | mmol m<sup>-2</sup> s<sup>-1</sup> |
| STOMATAL CONDUCTANCE | *g<sub>s</sub>* | mmol m<sup>-2</sup> s<sup>-1</sup> |
| CARBOXYLATION EFFICIENCY | *A/Ci* | - |
| MINIMUM LEAF CONDUCTANCE | *g<sub>min</sub>* | mmol m<sup>-2</sup> s<sup>-1</sup> |
| OPEN STATE QUANTUM EFFICIENCY | *Fv'/Fm'* | - |
| PHOTOCHEMICAL QUENCHING | *qP* | - |
| NON-PHOTOCHEMICAL QUENCHING | *qN* | - |

## Appendix

**Table A1.** Species-level averages with 95% confidence intervals for all measured morphological and physiological traits under control and drought treatments.

| Species | ID | Treatment | Group | B | AGB | BGB | RS | TLA | SLA | LDMC | GCL | SD |
| --- | --- | --- | --- | --- | --- | --- | --- | --- | --- | --- | --- | --- |
| *Bromus benekenii* | BB | Control | Graminoid | 0.105±0.011 | 0.062±0.007 | 0.042±0.005 | 0.7±0.07 | 17.41±1.68 | 326.48±37.85 | 0.17±0.01 | 50.73±2.74 | 58.55±3.6 |
| *Bromus benekenii* | BB | Drought | Graminoid | 0.052±0.007 | 0.029±0.004 | 0.023±0.004 | 0.81±0.14 | 6.01±1.17 | 233.96±21.41 | 0.27±0.05 | 41.07±2.39 | 51.19±4.38 |
| *Carex digitata* | CD | Control | Graminoid | 0.034±0.012 | 0.023±0.007 | 0.012±0.005 | 0.5±0.17 | 2.63±0.62 | 328.43±75.49 | 0.24±0.07 | NA | NA |
| *Carex digitata* | CD | Drought | Graminoid | 0.04±0.006 | 0.026±0.004 | 0.014±0.003 | 0.55±0.11 | 2.7±0.38 | 307.65±36.15 | 0.27±0.03 | NA | NA |
| *Geum urbanum* | GU | Control | Forb | 0.297±0.085 | 0.182±0.056 | 0.115±0.04 | 0.7±0.18 | 28.37±7.64 | 282.3±12.76 | 0.19±0.02 | 26.01±1.19 | 170.18±49.66 |
| *Geum urbanum* | GU | Drought | Forb | 0.153±0.031 | 0.101±0.019 | 0.052±0.017 | 0.54±0.23 | 18.49±3.62 | 224.87±15.28 | 0.31±0.05 | 23.57±1.99 | 172.74±54.72 |
| *Impatiens parviflora* | IP | Control | Forb | 0.587±0.098 | 0.396±0.069 | 0.192±0.05 | 0.51±0.12 | 67.44±11.73 | 580.32±80.94 | 0.06±0.01 | 20.4±1.56 | 186.93±67.38 |
| *Impatiens parviflora* | IP | Drought | Forb | 0.157±0.041 | 0.1±0.026 | 0.058±0.019 | 0.59±0.17 | 22.45±7.47 | 399.81±83.62 | 0.2±0.06 | 18.64±1.79 | 162.23±39.11 |
| *Lamium maculatum* | LM | Control | Forb | 0.961±0.196 | 0.566±0.112 | 0.395±0.116 | 0.73±0.21 | 49.12±7.24 | 335.53±32.16 | 0.32±0.04 | 25.95±1.85 | 216.19±44.63 |
| *Lamium maculatum* | LM | Drought | Forb | 0.271±0.074 | 0.189±0.062 | 0.082±0.036 | 0.47±0.15 | 16.93±7.46 | 246.63±25.8 | 0.36±0.09 | 22.71±1.75 | 203.55±44.54 |
| *Melica nutans* | MN | Control | Graminoid | 0.206±0.041 | 0.139±0.026 | 0.067±0.019 | 0.48±0.09 | 17.64±2.94 | 415.98±24.08 | 0.17±0.17 | 32.24±3.34 | 71.71±12.8 |
| *Melica nutans* | MN | Drought | Graminoid | 0.088±0.023 | 0.057±0.016 | 0.03±0.009 | 0.54±0.09 | 3.2±0.66 | 131.1±16.32 | 0.42±0.07 | 29.21±6.43 | 60.46±86.71 |
| *Poa nemoralis* | PN | Control | Graminoid | 0.366±0.08 | 0.185±0.031 | 0.181±0.057 | 0.96±0.25 | 8.61±0.78 | 388.79±34.37 | 0.03±0 | NA | NA |
| *Poa nemoralis* | PN | Drought | Graminoid | 0.062±0.013 | 0.035±0.009 | 0.027±0.009 | 0.85±0.32 | 2.25±0.57 | 254.33±64.81 | 0.14±0.06 | NA | NA |
| *Viola mirabilis* | VM | Control | Forb | 0.42±0.26 | 0.211±0.13 | 0.209±0.133 | 1.01±0.24 | 31.8±13.2 | 281.94±20.58 | 0.29±0.02 | 25.56±3.97 | 109.93±27.3 |
| *Viola mirabilis* | VM | Drought | Forb | 0.117±0.05 | 0.068±0.031 | 0.049±0.021 | 0.77±0.16 | 8.16±2.7 | 236.78±58.11 | 0.67±0.48 | 26.44±3.82 | 107.01±33.62 |
| *Veronica sublobata* | VS | Control | Forb | 0.368±0.056 | 0.349±0.056 | 0.019±0.002 | 0.06±0.01 | 39.34±5.49 | 598.2±67.11 | 0.09±0.02 | 27.85±2.06 | 116.95±30.4 |
| *Veronica sublobata* | VS | Drought | Forb | 0.076±0.012 | 0.07±0.011 | 0.007±0.002 | 0.1±0.03 | 13.43±3.49 | 252.89±64.69 | 0.25±0.13 | 23.13±1.85 | 134.35±31.84 |
| Species | ID | Treatment | Group | A | E | g<sub>s</sub> | iWUE | A/Ci | g<sub>min</sub> | F'v/F'm | qP | qN |
| *Bromus benekenii* | BB | Control | Graminoid | 8.21±0.59 | 2.7±0.24 | 79.23±7.98 | 0.104±0.005 | 0.039±0.003 | 9.47±1.35 | 0.61±0.02 | 0.37±0.03 | 0.75±0.03 |
| *Bromus benekenii* | BB | Drought | Graminoid | 1.21±0.77 | 0.43±0.2 | 11.68±5.56 | 0.092±0.024 | 0.006±0.004 | 7.96±0.72 | 0.45±0.04 | 0.27±0.03 | 0.9±0.03 |
| *Carex digitata* | CD | Control | Graminoid | 2.21±1.07 | 0.42±0.21 | 15.64±7.79 | 0.145±0.016 | 0.016±0.005 | 4.2±0.85 | 0.36±0.08 | 0.36±0.12 | 0.94±0.03 |
| *Carex digitata* | CD | Drought | Graminoid | 1.26±0.5 | 0.25±0.08 | 9.06±3.19 | 0.144±0.031 | 0.011±0.007 | 4.66±1.34 | 0.39±0.05 | 0.3±0.06 | 0.93±0.02 |
| *Geum urbanum* | GU | Control | Forb | 4.33±1.07 | 0.62±0.09 | 29.77±6.17 | 0.145±0.016 | 0.052±0.01 | 3.28±0.49 | 0.56±0.04 | 0.28±0.04 | 0.81±0.04 |
| *Geum urbanum* | GU | Drought | Forb | 0.33±0.34 | 0.07±0.04 | 2.3±1.47 | 0.094±0.415 | 0.002±0.002 | 3.71±1.43 | 0.51±0.02 | 0.17±0.06 | 0.86±0.02 |
| *Impatiens parviflora* | IP | Control | Forb | 4.83±0.74 | 0.85±0.15 | 34.17±6.09 | 0.144±0.005 | 0.032±0.005 | 8.78±2.79 | 0.81±0.01 | 0.19±0.02 | 0.05±0.06 |
| *Impatiens parviflora* | IP | Drought | Forb | 0.82±0.46 | 0.11±0.06 | 4.49±2.5 | 0.144±0.142 | 0.009±0.003 | 6.18±0.99 | 0.75±0.04 | 0.11±0.02 | 0.36±0.13 |
| *Lamium maculatum* | LM | Control | Forb | 5.31±0.97 | 1.5±0.29 | 46.61±9.01 | 0.115±0.007 | 0.027±0.005 | 2.93±0.6 | 0.54±0.01 | 0.28±0.04 | 0.83±0.02 |
| *Lamium maculatum* | LM | Drought | Forb | 0.11±0.11 | 0.18±0.03 | 4.88±0.88 | 0.023±0.02 | 0±0 | 1.76±0.87 | 0.53±0.02 | 0.09±0.02 | 0.85±0.02 |
| *Melica nutans* | MN | Control | Graminoid | 2.88±0.76 | 0.85±0.23 | 29.47±8.57 | 0.101±0.009 | 0.009±0.003 | 4.47±1.36 | 0.57±0.04 | 0.28±0.04 | 0.8±0.05 |
| *Melica nutans* | MN | Drought | Graminoid | NA | NA | NA | NA | NA | NA | NA | NA | NA |
| *Poa nemoralis* | PN | Control | Graminoid | 5.7±0.53 | 2.21±0.31 | 71.5±9.84 | 0.082±0.011 | 0.023±0.003 | 8.4±2.42 | 0.61±0.03 | 0.3±0.03 | 0.74±0.05 |
| *Poa nemoralis* | PN | Drought | Graminoid | 2.19±1.14 | 0.72±0.25 | 20.4±7.45 | 0.103±0.03 | 0.012±0.007 | 6.29±2.02 | 0.45±0.05 | 0.34±0.06 | 0.9±0.03 |
| *Viola mirabilis* | VM | Control | Forb | 7.55±1.25 | 3.62±1.02 | 106.22±32.01 | 0.075±0.018 | 0.029±0.005 | 4.46±1.27 | 0.59±0.01 | 0.27±0.04 | 0.77±0.01 |
| *Viola mirabilis* | VM | Drought | Forb | 0.44±0.63 | 0.42±0.25 | 11.09±6.53 | 0.024±0.028 | 0.002±0.002 | 4.29±1.63 | 0.51±0.09 | 0.14±0.03 | 0.85±0.08 |
| *Veronica sublobata* | VS | Control | Forb | 8.08±0.9 | 3.53±0.5 | 153.21±25.02 | 0.056±0.006 | 0.027±0.003 | 7.46±1.75 | 0.6±0.03 | 0.32±0.03 | 0.75±0.04 |
| *Veronica sublobata* | VS | Drought | Forb | 1.4±0.55 | 0.43±0.18 | 15.41±6.66 | 0.101±0.033 | 0.004±0.002 | NA | 0.43±0.07 | 0.27±0.11 | 0.9±0.04 |

**Table A2.** Results of Kruskal-Wallis for morphological traits and physiological traits with species, ecological guild, water regime treatment, and their interactions as fixed factors (p<0.05 *, p<0.01 **, p<0.001 ***).

| Factor | $\chi^2$ | df | p | Factor | $\chi^2$ | df | p |
| --- | --- | --- | --- | --- | --- | --- | --- |
| **B** |  |  |  | **A** |  |  |  |
| Species | 134.54 | 8 | *** | Species | 16.32 | 8 | * |
| Ecological Guild | 75.02 | 1 | *** | Guild | 0.26 | 1 | 0.61 |
| Treatment | 86.68 | 1 | *** | Treatment | 135.84 | 1 | *** |
| Species*Treatment | 229.43 | 17 | *** | Species*Treatment | 175.03 | 17 | *** |
| Guild*Treatment | 166.35 | 3 | *** | Guild*Treatment | 148.57 | 3 | *** |
| **AGB** |  |  |  | **E** |  |  |  |
| Species | 144.19 | 8 | *** | Species | 53.25 | 8 | *** |
| Guild | 92.53 | 1 | *** | Guild | 0.97 | 1 | 0.32 |
| Treatment | 76.66 | 1 | *** | Treatment | 113.29 | 1 | *** |
| Species*Treatment | 230.81 | 17 | *** | Species*Treatment | 180.64 | 17 | *** |
| Guild*Treatment | 174.88 | 3 | *** | Guild*Treatment | 124.69 | 3 | *** |
| **BGB** |  |  |  | **g<sub>s</sub>** |  |  |  |
| Species | 153.52 | 8 | *** | Species | 42.815 | 8 | *** |
| Guild | 14.07 | 1 | *** | Guild | 0.15 | 1 | 0.7 |
| Treatment | 56.88 | 1 | *** | Treatment | 126.06 | 1 | *** |
| Species*Treatment | 214.04 | 17 | *** | Species*Treatment | 181.15 | 17 | *** |
| Guild*Treatment | 72.31 | 3 | *** | Guild*Treatment | 135.39 | 3 | *** |
| **RS** |  |  |  | **WUEi** |  |  |  |
| Species | 125.24 | 8 | *** | Species | 70.146 | 8 | *** |
| Guild | 29.35 | 1 | *** | Guild | 6.07 | 1 | * |
| Treatment | 1.02 | 1 | 0.31 | Treatment | 7.67 | 1 | ** |
| Species*Treatment | 134.13 | 17 | *** | Species*Treatment | 120.36 | 17 | *** |
| Guild*Treatment | 30.73 | 3 | *** | Guild*Treatment | 25.362 | 3 | *** |
| **TLA** |  |  |  | **A/Ci** |  |  |  |
| Species | 144.48 | 8 | *** | Species | 5.38 | 8 | 0.72 |
| Guild | 114.03 | 1 | *** | Guild | 0.35 | 1 | 0.55 |
| Treatment | 64.03 | 1 | *** | Treatment | 128.31 | 1 | *** |
| Species*Treatment | 222.64 | 17 | *** | Species*Treatment | 164.64 | 17 | *** |
| Guild*Treatment | 183.7 | 3 | *** | Guild*Treatment | 144.69 | 3 | *** |
| **SLA** |  |  |  | **g<sub>min</sub>** |  |  |  |
| Species | 62.534 | 8 | *** | Species | 134.06 | 8 | *** |
| Guild | 3.84 | 1 | * | Guild | 33.47 | 1 | *** |
| Treatment | 72.45 | 1 | *** | Treatment | 2.81 | 1 | 0.09 |
| Species*Treatment | 165.97 | 17 | *** | Species*Treatment | 144.1 | 16 | *** |
| Guild*Treatment | 77.81 | 3 | *** | Guild*Treatment | 38.92 | 3 | *** |
| **LDMC** |  |  |  | **Fv/Fm** |  |  |  |
| Species | 121.14 | 8 | *** | Species | 104.33 | 8 | *** |
| Guild | 2.714 | 1 | 0.09 | Guild | 28.87 | 1 | *** |
| Treatment | 56.37 | 1 | *** | Treatment | 54.34 | 1 | *** |
| Species*Treatment | 190.6 | 17 | *** | Species*Treatment | 166.48 | 17 | *** |
| Guild*Treatment | 61.85 | 3 | *** | Guild*Treatment | 78.5 | 3 | *** |
| **GCL** |  |  |  | **qP** |  |  |  |
| Species | 145.98 | 8 | *** | Species | 78.314 | 8 | *** |
| Guild | 73.15 | 1 | *** | Guild | 41.702 | 1 | *** |
| Treatment | 5.91 | 1 | * | Treatment | 23.586 | 1 | *** |
| Species*Treatment | 155.95 | 16 | *** | Species*Treatment | 126.43 | 17 | *** |
| Guild*Treatment | 83.81 | 3 | *** | Guild*Treatment | 78.353 | 3 | *** |
| **SD** |  |  |  | **qN** |  |  |  |
| Species | 144.83 | 8 | *** | Species | 104.37 | 8 | *** |
| Guild | 100.06 | 1 | *** | Guild | 28.84 | 1 | *** |
| Treatment | 0.45 | 1 | 0.5 | Treatment | 54.39 | 1 | *** |
| Species*Treatment | 146.8 | 16 | *** | Species*Treatment | 166.58 | 17 | *** |
| Guild*Treatment | 103.7 | 3 | *** | Guild*Treatment | 78.55 | 3 | *** |

**Figure A1.**
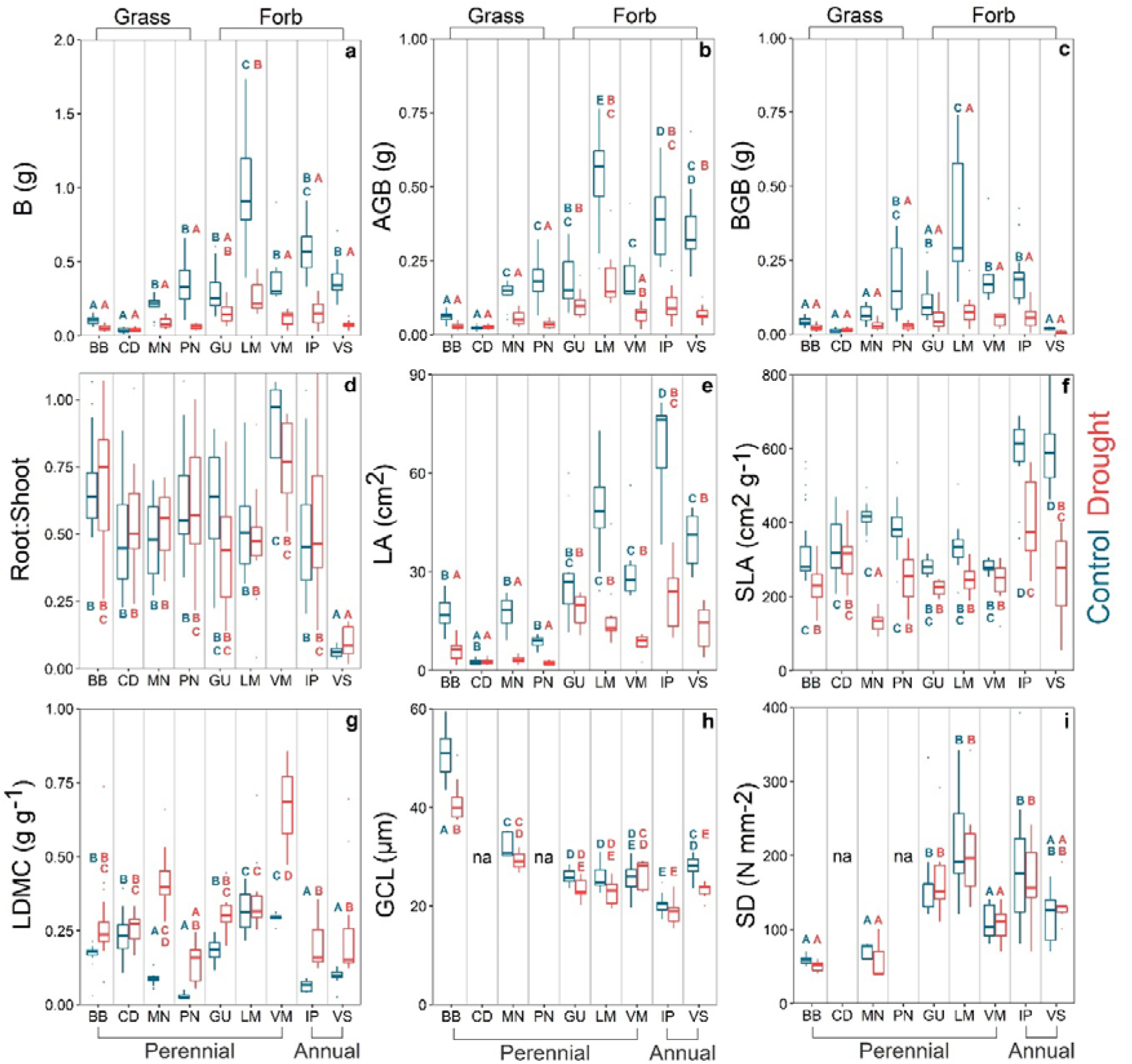
Species level boxplots of total biomass (a), aboveground biomass (b), belowground biomass (c), root to shoot ratio (d), total leaf area (e), specific leaf area (f), leaf dry matter content (g), stomatal guard cell length (h) and stomatal density (i). BB – *Bromus benekenii*, CD - *Carex digitata*, MN - *Melica nutants*, PN – *Poa nemoralis*, GU – *Geum urbanum*, LM – *Lamium maculatum*, VM – *Viola mirabilis*, IP – *Impatiens parviflora*, VS – *Veronica sublobata*.

**Figure A2.**
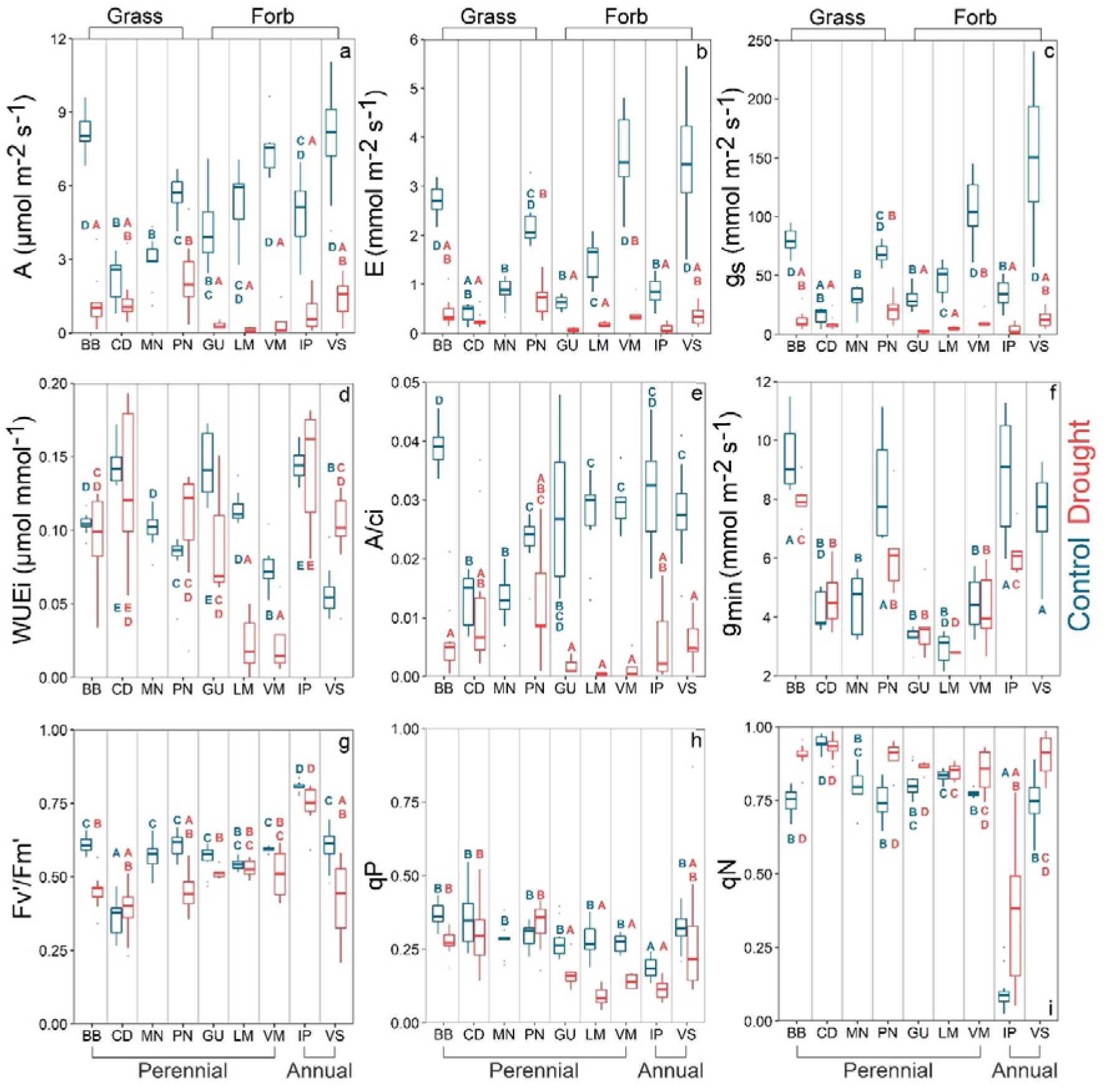
Species level boxplots of assimilation rate (a), transpiration rate (b), stomatal conductance (c), water use efficiency (d), carboxylation efficiency (e), minimal conductance (f), open state quantum efficiency (g), photochemical quenching (h), non-photochemical quenching (i). BB – *Bromus benekenii*, CD - *Carex digitata*, MN - *Melica nutants*, PN – *Poa nemoralis*, GU – *Geum urbanum*, LM – *Lamium maculatum*, VM – *Viola mirabilis*, IP – *Impatiens parviflora*, VS – *Veronica sublobata*.

**Figure A3.**
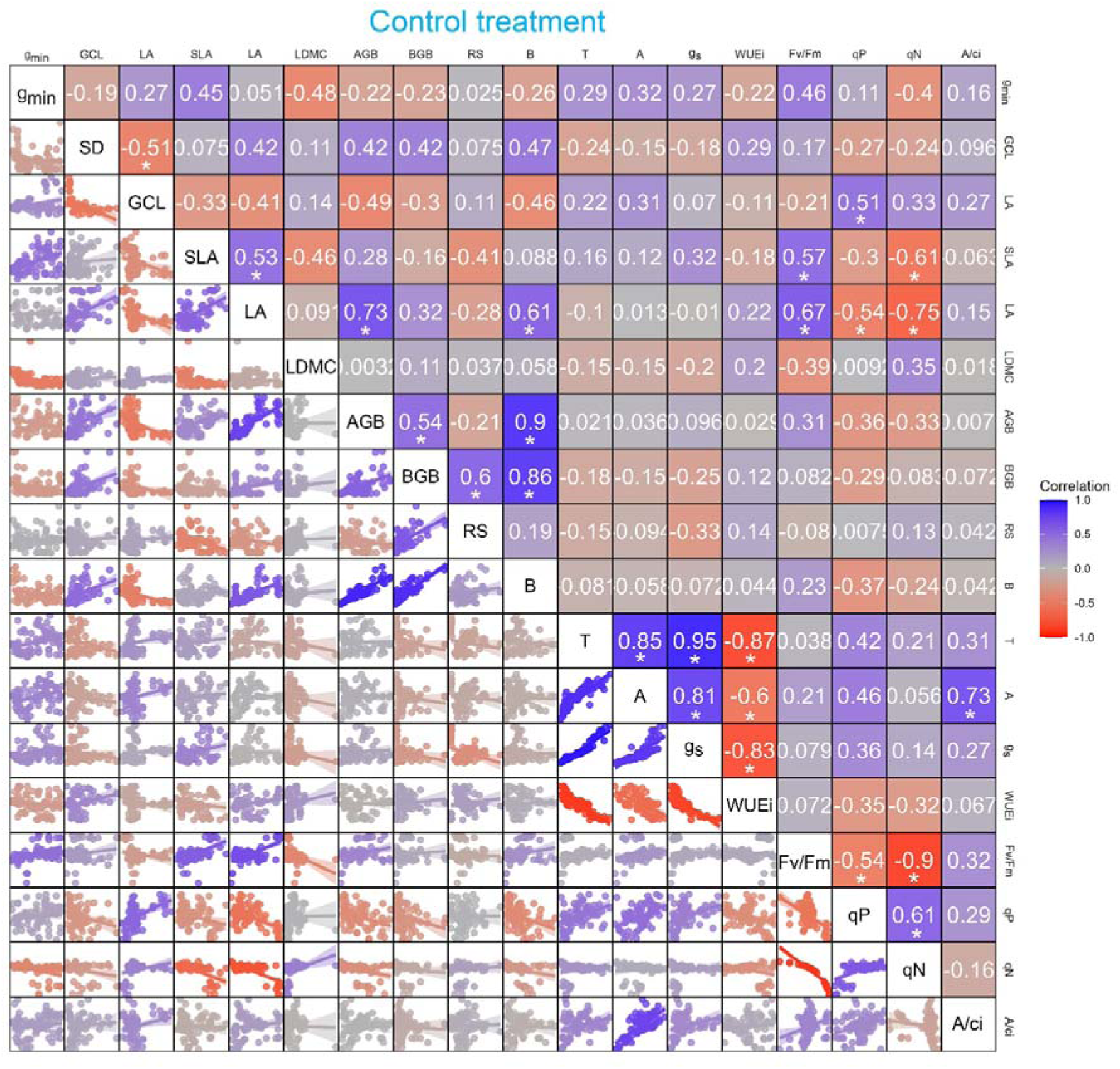
Pearson correlation matrix with linear trends for all evaluated traits under control treatment, asterisks mark significant correlations at P<0.05.

**Figure A4.**
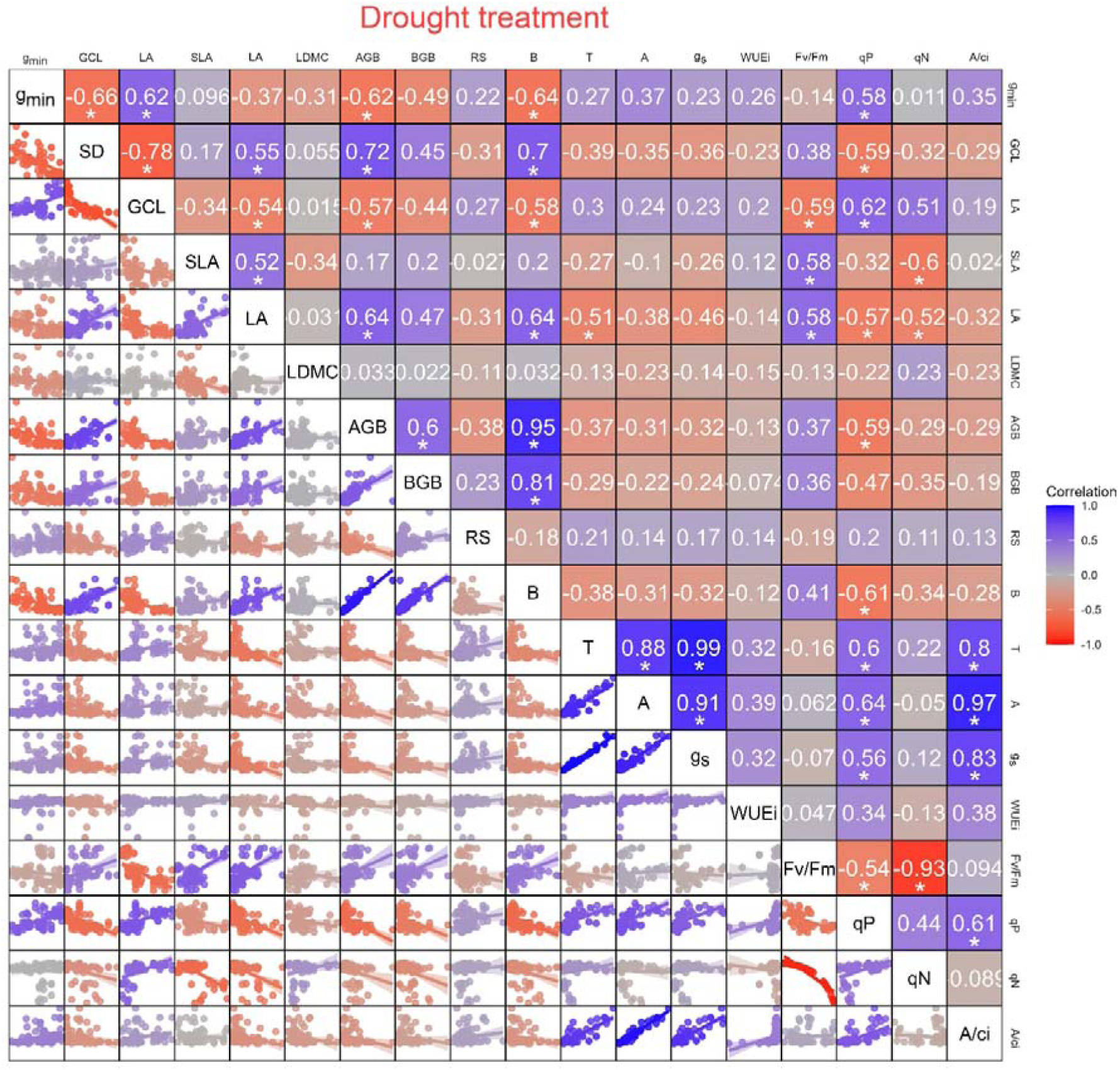
Pearson correlation matrix with linear trends for all evaluated traits under drought treatment, asterisks mark significant correlations at P<0.05.

